# K^+^ efflux through postsynaptic NMDA receptors suppresses local astrocytic glutamate uptake

**DOI:** 10.1101/2020.11.07.372722

**Authors:** Olga Tyurikova, Pei‐Yu Shih, Yulia Dembitskaya, Leonid P. Savtchenko, Thomas J. McHugh, Dmitri A. Rusakov, Alexey Semyanov

**Affiliations:** Brain Science Institute (BSI), RIKEN, Wako‐shi, Saitama, Japan; UCL Institute of Neurology, Queen Square, London, UK; Shemyakin‐Ovchinnikov Institute of Bioorganic Chemistry, Russian Academy of Sciences, Moscow, Russia; Sechenov First Moscow State Medical University, Moscow, Russia

## Abstract

Glutamatergic transmission prompts K^+^ efflux through postsynaptic NMDA receptors. K^+^ depolarizes local presynaptic terminals, boosting glutamate release, but whether it also affecting glutamate uptake remains unknown. Here, we find that the pharmacological blockade, or conditional knockout, of NMDA receptors suppresses the progressive use‐dependent increase in the amplitude and decay time of the astrocytic glutamate transporter current (I_GluT_), whereas blocking the astrocytic inward‐rectifying K^+^ channels prevents the decay time increase only. Glutamate spot‐uncaging reveals that local astrocyte depolarization, rather than extracellular K^+^ rises on their own, reduces the amplitude and prolong the decay of I_GluT_. Biophysical simulations confirm that local transient elevations of extracellular K^+^ can inhibit local glutamate uptake in fine astrocytic processes. Optical glutamate sensor imaging and a two‐pathway test relate postsynaptic K^+^ efflux to enhanced extrasynaptic glutamate signaling. Thus, postsynaptic K^+^ efflux facilitates presynaptic release while reducing local glutamate uptake.

## Introduction

K^+^ escapes brain cells through several mechanisms including K^+^ channels, transporters, and receptors. Postsynaptic α‐amino‐3‐hydroxy‐5‐methyl‐4‐isoxazolepropionic acid receptors (AMPARs) and NMDARs are both permeable for K^+^ (Wollmuth and Sobolevsky, 2004). K^+^ efflux through synaptic receptors makes synaptic transmission energetically costlier as it requires a larger number of Na^+^ ions to cross cell membranes to generate an excitatory postsynaptic current (EPSC). Nevertheless, K^+^ permeability of synaptic glutamate receptors has been evolutionary conserved, arguably reflecting a signaling role for K^+^. Transient accumulation of this ion in the narrow synaptic cleft depolarizes the adjacent presynaptic terminal, enhances action‐potential‐triggered presynaptic Ca^2+^ transients and therefore increases the probability of glutamate release (Contini et al., 2017; Hori and Takahashi, 2009). Patch‐clamp experiments combined with biophysical simulations have suggested that during synaptic transmission, the majority of K^+^ ions are released through postsynaptic NMDARs that remain activated for 100‐ 200 ms, as opposed to short‐lived K^+^ fluxes due to AMPAR activation or action potential repolarization phase (Shih et al., 2013). This retrograde signaling is excitatory activity‐dependent as K^+^ is released mostly through postsynaptic NMDARs which require postsynaptic depolarization to remove the Mg^2+^ channel block (Nowak et al., 1984). K^+^ released during synaptic transmission is cleared by extracellular diffusion, neuronal and astrocytic Na^+^/K^+^ ATPase, astrocytic inward‐rectifying K^+^ channels (K_ir_), and through various other channels and transporters. K^+^ current through inwardly rectifying astroglial channels (K_ir_) could last for hundreds of milliseconds in response to a single synaptic stimulus, suggesting a relatively long dwell time of extracellular K^+^ hotspots (Cheung et al., 2015; Lebedeva et al., 2018; Meeks and Mennerick, 2007). Because the membrane potential of astrocytes is predominantly determined by their K^+^ conductance, high extracellular K^+^ depolarizes the astroglial membrane, which can reduce voltage‐dependent glutamate uptake (Grewer et al., 2008; Grewer and Rauen, 2005; Mennerick et al., 1999). In addition to the membrane potential, astrocytic glutamate uptake depends on K^+^ and Na^+^ transmembrane gradients which are transiently reduced during synaptic transmission. How a combination of these factors affects extrasynaptic actions of glutamate, and therefore the excitatory signal spread in local circuitry, remains poorly understood.

## Results

### Activity‐dependent increase of IGluT amplitude and the decay time is mediated by postsynaptic NMDARs

Whole‐cell recordings were performed in CA1 *stratum radiatum* astrocytes in acute hippocampal slices of the mouse (Fig. 1A). Astrocytes were visually identified by their shape and the linear I‐V curve (Fig. 1B). A combined transporter (I_GluT_) and K^+^ (I_K_) current was evoked by extracellular stimulation of Schaffer collaterals (Fig. 1C). At the end of each experiment, 50 μM DL‐TBOA, a blocker of glutamate transporters, was added to isolate I_K_. The tail of I_K_ in the presence of DL‐TBOA was scaled to the tail of each individual combined current obtained in response to equivalent stimulation. The scaled I_K_ was subtracted from each corresponding trace of the combined current to obtain the time course of isolated I_GluT_ (Fig. 1C).

**Figure 1.**
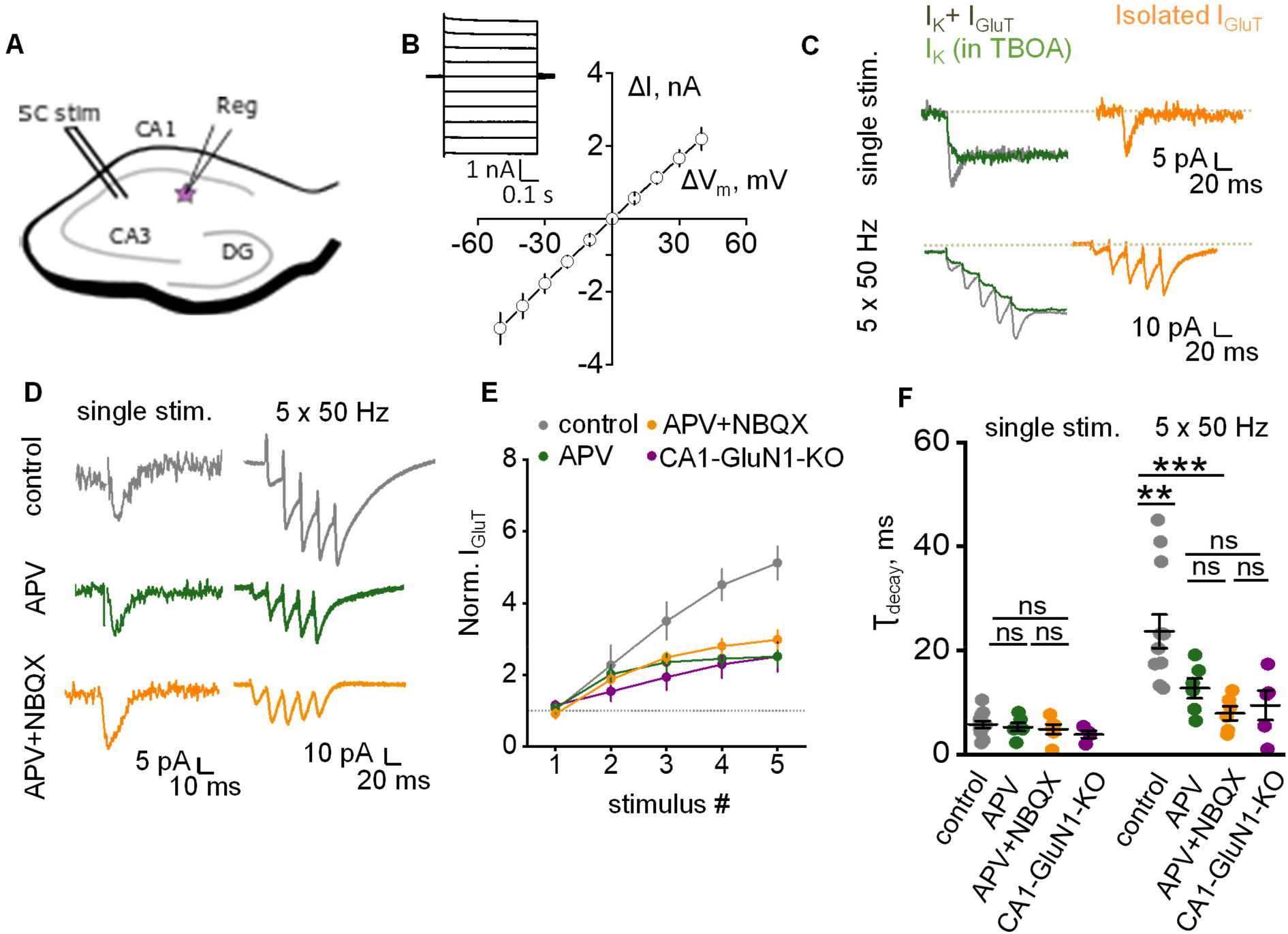
NMDAR‐mediated K^+^ efflux induces an activity‐dependent increase of I_GluT_ amplitude and decay time. **A.** A schematic showing stimulating (SC stim) and recording (Reg) electrode positions for the recording of synaptically‐induced currents in CA1 *stratum radiatum* astrocyte. **B**. Current‐voltage relationship of passive astrocyte. *Inset:* Astrocytic current (ΔI) was measured in response to voltage steps (ΔV_m_). **C.** Astrocytic currents induced by single and 5 × 50 Hz stimulation of Schaffer collaterals. DL‐TBOA application was used to isolate I_K_ (green), which was then subtracted from combined current (I_K_ + I_GluT_, grey) to obtain pure I_GluT_ (orange). **D.**Sample traces of I_GluT_ in response to single and 5 x 50 Hz stimulation in control (grey), in the presence of D‐APV (green) and in the presence of D‐APV + DL‐TBOA (orange). **E.** Amplitudes of I_GluT_ peaks during 5 × 50 Hz stimulation normalized to the amplitude of I_GluT_ in response to a single stimulus. Activity‐dependent facilitation in control (grey) was reduced by the application of D‐APV (green) or D‐APV + NBQX (orange) or in CA1‐GluN1‐KO mice (purple). **F.** τ_decay_ of I_GluT_ in response to single and 5 x 50 Hz stimulation. There was no significant difference in τ_decay_ of I_GluT_ in response to a single stimulus between control (grey), D‐APV (green), D‐APV + NBQX (orange) or in CA1‐GluN1‐KO mice (purple). τ_decay_ of I_GluT_ in response 5 x 50 Hz stimulation was significantly larger in control than in D‐APV, D‐APV + NBQX, or in CA1‐GluN1‐KO mice. The data are presented as mean ± SEM. ns p > 0.05, **p < 0.01 and ***p < 0.001, two‐sample *t*‐test.

Earlier work has suggested that NMDAR‐mediated K^+^ efflux produces activity‐dependent presynaptic depolarization and facilitation of glutamate release in synapses on CA1 pyramidal neurons (Shih et al., 2013). Indeed, we observed progressive facilitation of the I_GluT_ amplitude with 5 x 50 Hz stimulation; the facilitation was significantly reduced by 50 μM D‐APV, an NMDARs antagonist (*p*_APV_ < 0.001; *p*_stimulus_ < 0.001; *p*_APV*stimulus_ = 0.01; *n =* 6; two‐way RM ANOVA; Fig. 1D, E). The residual I_GluT_ facilitation can be attributed to an increase in presynaptic release probability due to mechanisms independent of the NMDARs‐mediated K^+^ efflux, such as the accumulation of residual Ca^2+^ in the presynaptic terminal. Application of 25 μM NBQX, an AMPAR antagonist, in addition to D‐APV, did not increase the effect, suggesting a minor role for AMPARs in the regulation of glutamate release. Activity‐dependent facilitation of I_GluT_ in the mice with conditional knockout of the GluN1 subunit of NMDARs in CA1 pyramidal neurons (CA1‐GluN1‐KO) (McHugh et al., 1996; Wu et al., 2012) was similar to the effect of D‐APV, confirming the involvement of postsynaptic NMDARs.

The decay time constant (τ_decay_) of I_GluT_ recorded in response to a single stimulus was not significantly affected by D‐APV or by co‐application of D‐APV and NBQX (control: 5.75 ± 0.65 ms, *n =* 12; D‐APV: 5.26 ± 0.80 ms, *n =* 6; *p =* 0.64 for the difference between control and D‐APV; D‐APV + NBQX: 4.83 ± 0.92 ms, *n =* 6; *p =* 0.72 for the difference between control and D‐APV + NBQX; two‐sample *t*‐test; Fig. 1F). The τ_decay_ of I_GluT_ recorded in response to a single stimulus in CA1‐GluN1‐KO mice was also not significantly different from τ_decay_ in control mice (KO mice: 3.82 ± 0.69 ms, *n =* 4; *p =* 0.07 for the difference with control mice; two‐sample *t*‐test; Fig. 1F).

The values of τ_decay_ became significantly larger in response to 5 × 50 Hz stimulation (23.65 ± 3.22ms, *n =* 12; *p <* 0.001 for the difference with single stimulus; paired‐sample *t*‐test; Fig. 1F). This increase was significantly reduced by bath application of D‐APV (τ_decay_ = 12.71 ± 1.91 ms, *n =* 6, *p =* 0.01 for the difference with control, two‐sample *t*‐test; Fig. 1F). Adding NBQX to D‐APV produced a further reduction in τ_decay_, albeit statistically insignificant (τ_decay_ = 7.91 ± 1.35 ms, *n =* 6, *p <* 0.001 compared to control, *p =* 0.13 compared to D‐APV, two‐sample *t*‐test; Fig. 1E). These results are consistent with an earlier suggestion that synaptically released glutamate does not overwhelm glutamate transporters upon the blockade of AMPARs and NMDARs (Diamond and Jahr, 2000). τ_decay_ in CA1‐GluN1‐KO mice was not significantly different from τ_decay_ in D‐APV (KO mice: 9.42 ± 2.82 ms, *n =* 5; *p =* 0.21 for the difference with D‐APV, two‐sample *t*‐test; Fig. 1F). These findings suggest that postsynaptic NMDARs are required both for the activity‐dependent facilitation of glutamate release and for the activity‐dependent reduction of glutamate uptake.

### Activity‐dependent increase of I_GluT_ decay time does not depend on afferent recruitment

NMDARs activation requires voltage‐dependent unblocking of the receptor channel (Nowak et al., 1984). Therefore, greater postsynaptic depolarization should, in theory, recruit more NMDARs and thus produce larger K^+^ efflux. To test this hypothesis, we recorded I_GluT_s at two different stimulation strengths (Fig. 2A). Initially, the stimulation strength was adjusted to achieve I_GluT_ ~ 10 pA (weak stimulus), then the strength was doubled (strong stimulus). Strong stimulation increased the I_GluT_ to 17.07 ± 1.8 pA (*n =* 8; *p =* 0.03, paired sample *t*‐test; Fig. 2A). Surprisingly, we did not observe a significant enhancement in the activity‐dependent facilitation of I_GluT_ upon stronger stimulation (*p*_strengh_ = 0.41; *p*_stimulus_ < 0.001; *p*_strengh*stimulus_ = 0.99; *n =* 7; two‐way RM ANOVA; Fig. 2B). Nor did we observe a significant difference in τ_decay_ between weak and strong stimulation groups, either with one or with five stimuli tests (single stimulus: 5.57 ± 1.3 ms for weak stimulation, 5.89± 1.34 ms for strong stimulation, *p =* 0.87, paired sample *t*‐test; 5 stims: 27.24 ± 5.29 ms for weak stimulation, 25.29 ± 3.77 ms for strong stimulation, *p =* 0.78, paired sample *t*‐test; *n =* 6; Fig. 2C). Thus, although stronger stimulation releases more glutamate per tissue volume, hence generates larger uptake currents, the current kinetics appears unaffected. This is likely because glutamate transporter binding and uptake occur only inside the microscopic vicinity of individual synapses, within 5 ‐ 10 ms post‐release. Unless such ‘uptake hotspots’ substantially overlap in the tissue volume, engaging additional synapses would not be expected to affect glutamate uptake kinetics.

**Figure 2.**
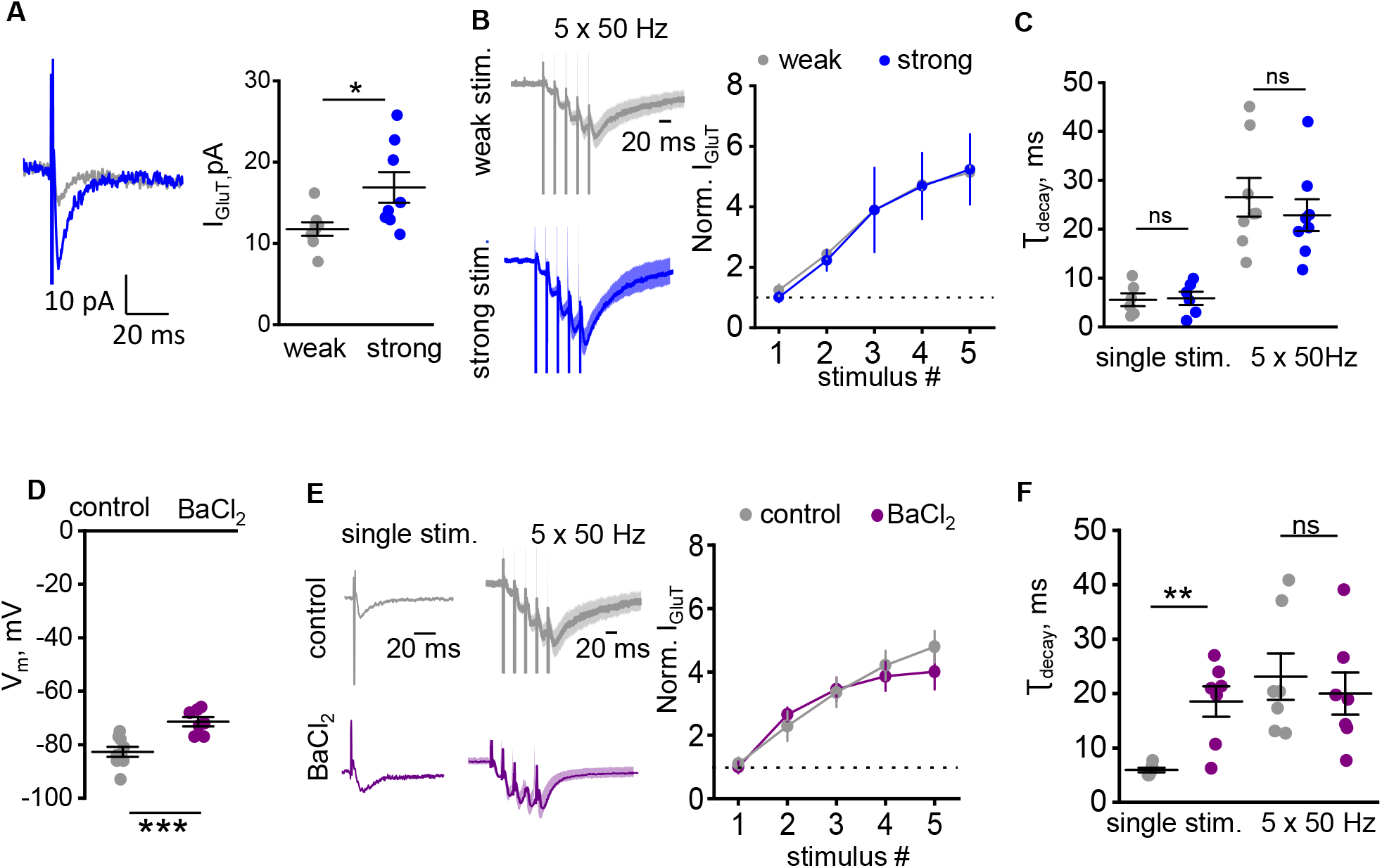
Activity‐dependent increase of I_GluT_ decay time does not depend on the stimulus strength but abolished by the blockade of K_ir_. **A.** *Left:* Traces of I_GluT_ recorded with weak (grey) and strong (weak*2; blue) single stimulation of Shaffer collaterals. *Right:* The summary plot showing an increase in the amplitude of I_GluT_ by increasing the stimulation strength twice (from weak to strong). **B.** The increase in stimulation strength did not affect the activity‐dependent facilitation of I_GluT_. *Left:* The traces of I_GluT_ in response to 5 x 50 Hz stimulation normalized to the amplitude of I_GluT_ in response to single weak stimulation. The traces were averaged in all experiments and presented as mean ± SEM. *Right:* The summary plot of normalized I_GluT_ amplitude in response to each stimulus in the train. **C.** τ_decay_ of I_GluT_ in response to single and 5 x 50 Hz stimulation was not affected by the increase in stimulus strength. Grey circles – weak stimulation; blue circles – strong stimulation. **D.** The summary graph showing the effect of BaCl_2_ astrocyte membrane potential. **E.** *Left:* Normalised traces of I_GluT_ recorded with control intracellular solution (grey) and the solution containing K_ir_ channels blocker ‐ BaCl_2_ (purple). The traces for 5 × 50 Hz stimulation were averaged in all experiments and presented as mean ± SEM. *Right:* The summary plot of normalized I_GluT_ amplitude in response to each stimulus in the train. **F.** The summary graph showing the effect of BaCl_2_ on τ_decay_ of I_GluT_ in response to single and 5 × 50 Hz stimulation. Grey circles – control; purple circles – BaCl_2_. The data are presented as mean ± SEM. ns p > 0.05, *p < 0.05 and **p < 0.01, paired‐sample (A, D) and two‐sample (C,F) *t*‐test.

To provide further controls, we have carried out experiments in which astrocytic K^+^ currents (I_K_) were compared directly under strong and weak stimuli, in conditions of single and burst stimuli. As expected, I_K_ increased with the increased stimulus strength or number. Importantly, in response to a stimulus burst, the use‐dependent increase in I_K_ was similar under either weak or strong stimuli (Fig. S1), thus arguing that it scales linearly independently of the number of synapses/afferent fibers involved.

### Activity‐dependent increase of I_GluT_ decay time is abolished by the blockade of astrocytic K_ir_

Next, we asked if the activity‐dependent increase in τ_decay_ involves the K_ir_‐dependent depolarization of astrocytic leaflets. Earlier work has shown that dialysis of astrocytes with 100 μM BaCl_2_ blocks K_ir_ channels responsible for I_K_ in astrocytes (Afzalov et al., 2013). Here, BaCl_2_ produced a small but significant depolarization of the recorded astroglia (V_m_ = ‐82.66 ± 1.85 in control, *n =* 9; V_m_= ‐71.42 ± 1.73 in control, *n =* 7; *p <* 0.001 two‐sample *t*‐test; Fig. 2D). Surprisingly, BaCl_2_ had no significant effect on I_GluT_ facilitation (*p*_BaCl2_ = 0.26; *p*_stimulus_ < 0.001; *p*_BaCl2*stimulus_ = 0.34; *n =* 6; two‐way RM ANOVA; Fig. 2E). This likely explanation is that K^+^ dynamics in the synaptic cleft depends mainly on diffusion escape rather than on the K_ir_‐ mediated clearance (Meeks and Mennerick, 2007). On the other hand, BaCl_2_ significantly increased τ_decay_ in response to a single stimulus (5.93 ± 0.41 ms, *n =* 6, *p =* 0.01 for the difference with control, two‐sample *t*‐test; Fig. 2F). This finding is consistent with the reduction of electrogenic glutamate uptake following astrocyte depolarization that occurs in perisynaptic astrocytic leaflets upon blockade of K_ir_. Indeed, astrocytes have very leaky membranes in which voltage clamp can be easily disturbed by local currents (e.g., (Savtchenko et al., 2018)). No further activity‐dependent prolongation changes in τ_decay_ were observed (τ_decay_ in response to 5 x 50 Hz stimulation = 20.03 ± 4.59 ms, *n =* 6, *p =* 0.02, paired‐sample *t*‐ test; Fig. 2F). Notably, τ_decay_ in these tests was not significantly different from τ_decay_ in response to 5 x 50 Hz stimulation in control conditions (*p =* 0.61, two‐sample *t*‐test; Fig. 2F). These data suggest that the K_ir_‐dependent membrane depolarization, but not a decrease in the transmembrane K^+^ gradient, is responsible for the activity‐dependent prolongation of I_GluT_ in astrocytes.

### Astrocyte depolarization but not extracellular K^+^ affects I_GluT_

Our findings suggest that K^+^ efflux through postsynaptic NMDARs decreases glutamate uptake. The two candidate underlying mechanisms are (1) depolarization of the astrocytic membrane and (2) a decrease in the astrocytic transmembrane K^+^ gradient. Synaptically released glutamate rapidly binds to astrocytic transporters (Diamond and Jahr, 1997). From the bound state, glutamate^‐^ can be translocated into the astrocyte cytoplasm, along with 3 Na^+^ and 1 H^+^, in exchange for 1 K^+^ (Grewer and Rauen, 2005). Therefore, the glutamate translocation step is both K^+^ ‐ and voltage‐dependent (Grewer et al., 2008; Mennerick et al., 1999). To assess the relative contributions of K^+^ and voltage, we recorded I_GluT_ induced by the single‐pulse spot uncaging of glutamate that generates a typical single‐synapse (unitary) EPSC (uI_GluT_), as described previously (Henneberger et al., 2020), near the astrocyte soma, to minimize voltage‐clamp error (Fig. 3A), under the two sets of conditions as follows. Firstly, we obtained the relationship between the extracellular K^+^ concentration and astrocyte membrane potential (Fig. 3B). An increase in extracellular K^+^ produced similar astrocyte depolarization as previously reported (Ge and Duan, 2007). Second, we mimicked the K^+^‐induced depolarization by holding the cell in voltage‐clamp mode at the membrane potentials seen at high K^+^. The membrane depolarization alone reduced uI_GluT_ to the same degree as did the corresponding K^+^ concentration uI_GluT_ (*p*_K+_ = 0.18; *p*_depolarization_< 0.001; *p*_K+*depolarization*_ = 0.59; *n =* 7; two‐way RM ANOVA; Fig. 3C). These results suggest that depolarization alone can attenuate glutamate uptake, thus occluding the effects of K^+^ elevations. We also observed a depolarization‐dependent increase in τ_decay_ of uI_GluT_ (*p* = 0.007, *n* = 8, one‐way RM ANOVA; Fig. 3D). Correspondingly, when different extracellular K^+^ concentrations were applied under a constant membrane potential of ‐85 mV, no significant change in the amplitude of uI_GluT_ (*p* = 0.14, *n* = 4; 8; 4, one‐way RM ANOVA) or in τ_decay_ (*p* = 0.003, *n* = 11; 7; 4 for each condition one‐way RM ANOVA) was observed (Fig. 3E,F). These findings suggest that physiologically relevant elevations in extracellular K^+^ affect I_GluT_ through astrocyte depolarization rather than by reducing the driving force for K^+^.

**Figure 3.**
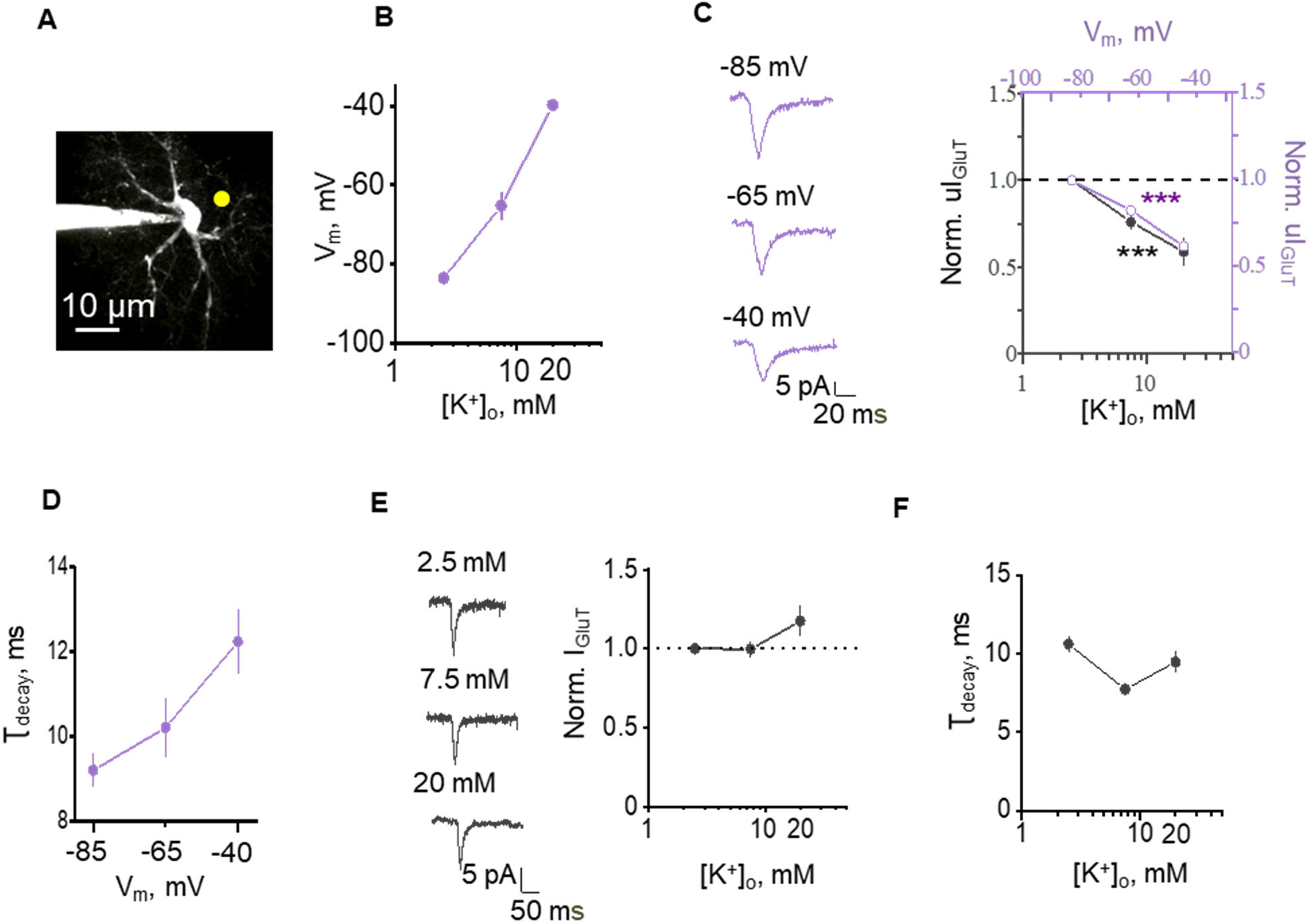
Depolarization of astrocyte rather than a decrease in K^+^ gradient suppresses glutamate uptake. **A.** An image of astrocyte filled with Alexa Fluor 594. An uncaging spot was in proximity to soma to ensure proper voltage clamp. **B.** Astrocyte depolarization produced by an increase in [K^+^]_o_. **C.** *Left*, sample traces of uI_GluT_ produced by glutamate uncaging while holding astrocyte at different membrane potential (V_m_). *Right*, the summary plot showing normalized uI_GluT_ amplitude in the cells voltage‐clamped at −85, −65, and −40 mV (purple circles) and in the cells clamped at the same voltages but with [K^+^]_o_ corresponding to each voltage as in B (2.5, 7.5, and 20 mM; black circles). K^+^ elevation did not have any additional effect on the cell depolarization. **D.** The summary plot showing τ_decay_ of uI_GluT_ in the cells voltage‐clamped at ‐85, ‐65, and ‐40 mV. **E.** *Left*, sample traces of uI_GluT_ produced by glutamate uncaging while holding astrocyte at fixed V_m_ = −85 mV but with different [K^+^]_o_ = 2.5, 7.5, and 20 mM; *Right*, the summary plot showing normalised uI_GluT_ amplitude in the cell exposed to different [K^+^]_o_. **F.** The summary plot showing τ_decay_ of uI_GluT_ in the cells voltage‐clamped at −85 mV and exposed to [K^+^]_o_ = 2.5, 7.5, and 20 mM. The data are presented as mean ± SEM. ***p < 0.01, one‐sample *t*‐test.

### A biophysical underpinning of the relationship between extracellular K^+^ and glutamate uptake

To understand through which biophysical mechanism the extracellular K+ rises can affect the kinetics of astrocytic glutamate uptake, we employed a realistic, multi‐compartmental 3D model of a *stratum radiatum* astrocyte, which was developed and validated earlier using the NEURON‐compatible model builder ASTRO (Savtchenko et al., 2018). The model features known membrane astrocytic mechanisms, including GLT‐1 transporters and K_ir_4.1 K^+^ channels, distributed in the model membrane to match multiple experimental observations (Materials and Methods).

First, we employed the model to simulate extracellular K^+^ elevation (from 2.5 to 5 mM over a 20 μm spherical area, 1 s duration) within the 3D astrocyte territory (Fig. 4A). This generated local K^+^ influx through K_ir_4.1 channels, triggering diffuse equilibration of intracellular K^+^ across the tortuous cell lumen (Fig. 4B). The evolving dynamics of extra‐ and intracellular K^+^ was paralleled by local membrane depolarization (Fig. 4C). Aiming to mimic synchronous multi‐synaptic glutamate release, we also simulated a brief extracellular glutamate rise (0.1 mM for 1 ms, at 0.9 s post‐onset)within a 3 μm spherical area (Fig. 4D) placed inside the area of extracellular K^+^ elevation. To understand the effect of extracellular K^+^ rises on glutamate uptake, we, therefore, ran the glutamate release test in two conditions, one at the baseline extracellular K^+^ and one during K^+^ elevation, as above. In these tests, cloud‐computing modeling could reveal 3D membrane profiles of transporter current at high (nanoscopic) resolution (Fig. 4E). The I_GluT_ value volume‐averaged over the same glutamate uptake hotspot showed a significantly lower (by ~15%) amplitude during K^+^ elevation (Fig. 4F). This decrease was similar to the effect of the K^+^ application of I_GluT_ induced by glutamate uncaging (Fig 3C). These simulation results provide a theoretical biophysical basis to the hypothesis that transient elevations of extracellular K^+^ could inhibit local glutamate uptake by astrocytes.

**Figure 4.**
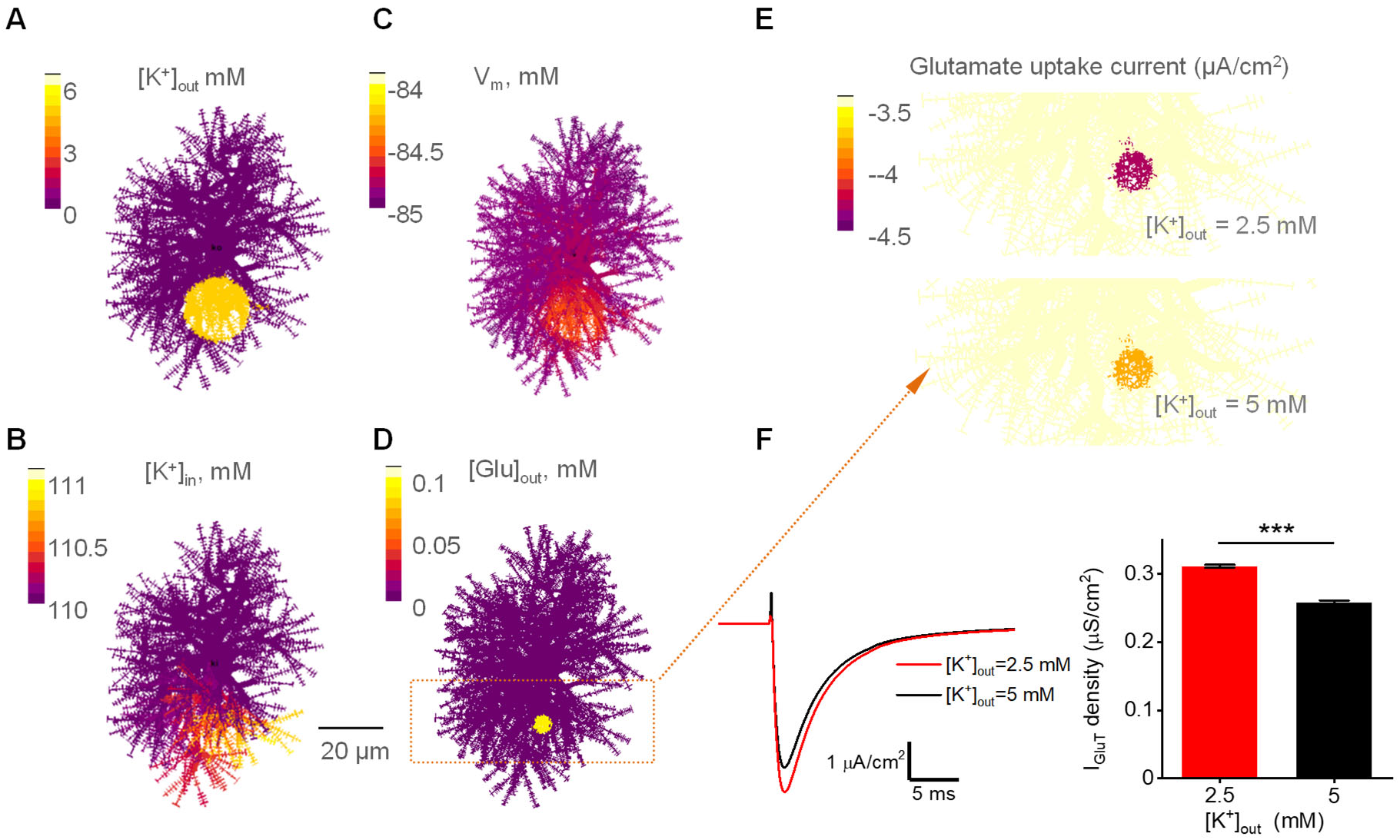
Realistic astrocyte model reveals the effect of extracellular K^+^ rises on local dynamics of astrocytic glutamate uptake. **A‐C,** Cell‐shape diagram illustrating a realistic 3D ASTRO‐NEURON‐based model of a CA1 astrocyte that incorporates known astrocytic membrane mechanisms including GLT‐1 transporters and K_ir_4.1 channels. Colour‐coding provides snapshots (2D projections, 0.9 s post‐onset), showing how a local [K^+^]_out_ rising (from 2.5 to 5 mM over a 20 μm spherical area, 1 s duration) affects [K^+^]_in_ (C) and transmembrane voltage (A_3_). **D,** Cell‐shape diagram illustrating ‘multi‐synaptic’ release of glutamate (0.1 mM for 1 ms, at 0.9 s post‐onset) within a small extracellular area (colour hotspot, false colour scale applies). **E,**Landscape of glutamate uptake current density (1 ms post‐release, overall ROI shown as a dotted rectangle in B), under baseline [K^+^]_out_ = 2.5 mM (*top*) and during [K^+^]_out_ elevation to 5 mM as shown in A‐C. **F,** *Left,* Glutamate uptake current kinetics over the affected cell area (colour hotspot) shown in C, at two [K^+^]_out_, as indicated. *Right,* Summary of a series of simulation tests recording I_GluT_ density at ten different distances from the modelled cell soma. mean ± SEM, n = 10, ***p < 0.001 paired‐sample *t*‐test.

### Synaptic K^+^ efflux boosts glutamate spillover

Changes in I_GluT_ may not faithfully reflect glutamate concentration changes in the extracellular space as I_GluT_ reflects, in large part, the rate of glutamate translocation across the astrocytic membrane. Glutamate binding occurs when free transporters are in abundance. However, during repetitive stimuli, slowing down or reducing I_GluT_ would reflect a greater fraction of transporters bound by glutamate (before the transmembrane transfer step) on the astrocytic surface. As the fraction of free transporters on the astrocytic surface decreases, the probability of glutamate molecules travelling further from the release sites increases. In this case, I_GluT_ would reflect the dynamics of glutamate escape and removal to a much greater degree. When the glutamate translocation rate is reduced by astrocytic depolarization, more transporters can buffer extracellular glutamate (Diamond and Jahr, 1997). Such increased glutamate buffering (binding‐unbinding) by transporter molecules could increase the dwell time of glutamate near the synaptic cleft (Lehre and Rusakov, 2002; Zheng et al., 2008). Therefore, we attempted to assess the dynamics of extracellular glutamate concentration using the tests as follows. First, we recorded excitatory postsynaptic potentials (EPSPs) in CA1 pyramidal neurons in response to a single stimulus and to 5 × 50 Hz stimulation, in the presence of 100 μM picrotoxin, a GABA_A_ receptor antagonist (Fig. 5A). The cut was made between CA1 and CA3 regions to prevent epileptiform activity. One of the two intracellular solutions was used: potassium methanesulfonate (KMS)‐based or N‐methyl‐D‐glucamine (NMDG)‐based. Having NMDG in the postsynaptic cell reduced the use‐dependent facilitation of EPSPs (Fig. 5A), lending support to the earlier finding that K^+^ efflux boosts presynaptic release efficacy (Shih et al., 2013). In contrast to NMDG‐based solution, KMS‐based solution allowed K^+^ efflux through postsynaptic receptors, which curtailed EPSP. Therefore, the decay time constant (τ_decay_) of EPSP was smaller in KMS than in NMDG recordings (KMS: 91.6 ± 20.63 ms, *n =* 7; NMDG: 210.8 ± 24.5 ms, *n =* 6; *p =* 0.003, two‐sample *t*‐test; Fig. 5B). The τ_decay_ of burst EPSP in response to 5 × 50 Hz stimulation was increased in KMS but not in NMDG (KMS: τ_decay_ = 136.52 ± 17.02 ms, τ_burst/single_ = 1.4 ± 0.14, *n =* 6, *p =* 0.03, single‐sample *t*‐test for the ratio; NMDG: τ_decay_ = 221.9 ± 23.5, τ_burst/single_ = 1.06 ± 0.05, *n =* 6, *p =* 0.34, single‐sample *t*‐test for the ratio; *p < 0.001*, two‐sample *t*‐test between ratios; Fig. 5C, D).

**Figure 5.**
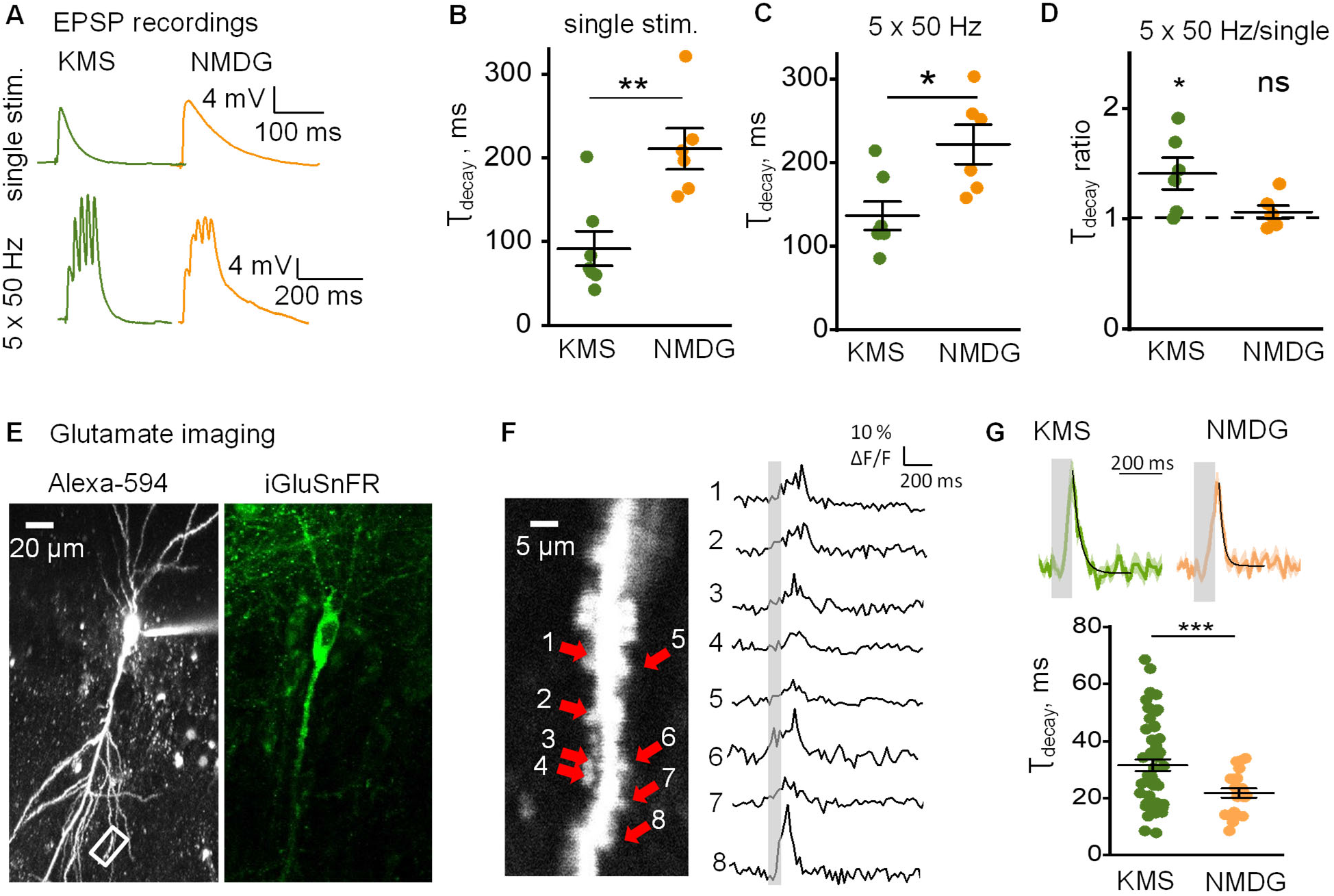
Synaptic K^+^ efflux increases glutamate dwell time in the synaptic cleft and enhances its spillover. **A.** Sample traces of EPSPs recorded in CA1 pyramidal neurons filled with either KMS- or NMDG‐based internal solutions in response to single‐pulse or 5 x 50 Hz stimulation, as indicated. **B.** and **C.** The summary plot of τ_decay_ of EPSPs in response to a single‐pulse (B) and 5 × 50 Hz (C) stimulation. Replacement of intracellular K^+^ (green) for NMDG (orange) significantly increased τ_decay_ in both cases. **D.** The summary plot showing the ratio of τ_decay_ of EPSP in response to 5 × 50 Hz stimulation to τ_decay_ of EPSP in response to single‐pulse stimulation. The activity‐dependent prolongation of EPSP was observed when the cell was filled with KMS‐ but not with an NMDG‐based solution. **E.** CA1 pyramidal neuron expressing iGluSnFR loaded with fluorescent dye Alexa–594 via patch pipette, shown in two emission channels, as indicated. **F.** *Left*, zoomed‐in area boxed in E; arrows, analyzed dendritic spines. *Right*, glutamate traces recorded at the corresponding spines. Grey bar ‐ stimulation. **G.** Averaged traces and a summary plot showing the τ_decay_ of glutamate transients recorded from 53 dendritic spines (5 cells) and 20 dendritic spines (3 cells) with KMS and NMDG‐based solutions, respectively. The substitution of K^+^ with NMDG shortened the glutamate dwell time around the synapses. The data are presented as mean ± SEM. ns p > 0.05, *p < 0.05, **p < 0.01 and ***p < 0.001, two‐sample (B,C,G) and one‐sample (D) *t*‐test.

This finding suggests that glutamate spillover is regulated by K^+^ efflux through postsynaptic receptors. Alternatively, this result may reflect the voltage and activity‐dependent regulation of K^+^ current that curtails EPSPs (Ichinose et al., 2003). Therefore, we next attempted to directly evaluate extracellular glutamate transient with genetically encoded glutamate sensor iGluSnFR (Fig. 5E). CA1 pyramidal neurons expressing the sensor (methods as described earlier (Jensen et al., 2019; Jensen et al., 2017)) were loaded through the patch pipette with either KMS‐ or NMDG‐based intracellular solution. We documented synaptically evoked extracellular glutamate transients using 500 Hz line‐scans placed across visually identified dendritic spines in the second‐order apical dendrite branches (Fig. 5F). Glutamate responses to burst stimulation (5 x 50 Hz) were thus recorded. The value of τ_decay_ for glutamate transients was significantly larger if the cell was loaded with the KMS‐based intracellular solution compared to the NMDG‐based solution (KMS: 31.56 ± 2.04 ms, *n =* 53 spines from 5 cells; NMDG: 21.8 ± 1.65 ms, *n =* 20 spines from 3 cells; *p <* 0.001, two‐sample *t*‐test; Fig. 5G). This finding supports the suggestion that activity‐dependent prolongation of EPSPs is due to glutamate spillover boosted by K^+^ efflux from the postsynaptic terminal.

### Synaptic K^+^ efflux boosts intersynaptic crosstalk

Enhanced glutamate spillover can facilitate intersynaptic crosstalk via high‐affinity NMDARs. To test if synaptic K^+^ efflux promotes the crosstalk, we performed a modified two‐pathway experiment (Henneberger et al., 2020; Scimemi et al., 2004). Briefly, NMDARs‐mediated EPSPs in CA1 pyramidal neurons were pharmacologically isolated with NBQX and picrotoxin, AMPAR and GABA_A_ receptor blockers, and recorded during depolarizing voltage steps to −40 mV. Two bipolar electrodes were placed in CA1 *stratum radiatum* on the opposite sides of the slice to recruit independent afferent pathways of Schaffer collaterals (Fig. 6A). The lack of cross‐facilitation of EPSPs in response to paired stimulation (first‐second and second‐first pathway) confirmed pathways independence (Scimemi et al., 2004) (Fig. 6B). After baseline recordings of EPSPs in both pathways, the stimulation of one pathway was stopped (silent pathway), and 4 μm MK‐801, an activity‐dependent NMDARs blocker, was applied. High‐frequency stimulation (HFS) was delivered to the second pathway (active pathway). This stimulation led to the relatively rapid blockade of NMDARs at the active pathway synapses and some synapses at the silent pathway, which were reached by the glutamate escaping the active synapses (Fig. 6C). The proportion of synapses blocked by MK‐801 at the silent pathway was estimated upon the drug washout and was significantly larger when the cell was filled with KMS‐ than NMDG‐based intracellular solution (EPSP: 42 ± 4 % of baseline for KMS‐based solution, *n =* 7; 63.9 ± 6 % of baseline for NMDG‐based solution, *n =* 6; *p =* 0.01, two‐sample *t*‐test; Fig. 6C,D). This finding suggests that K^+^ efflux through postsynaptic receptors promotes glutamate spillover‐mediated intersynaptic crosstalk involving high‐affinity NMDARs.

**Figure 6.**
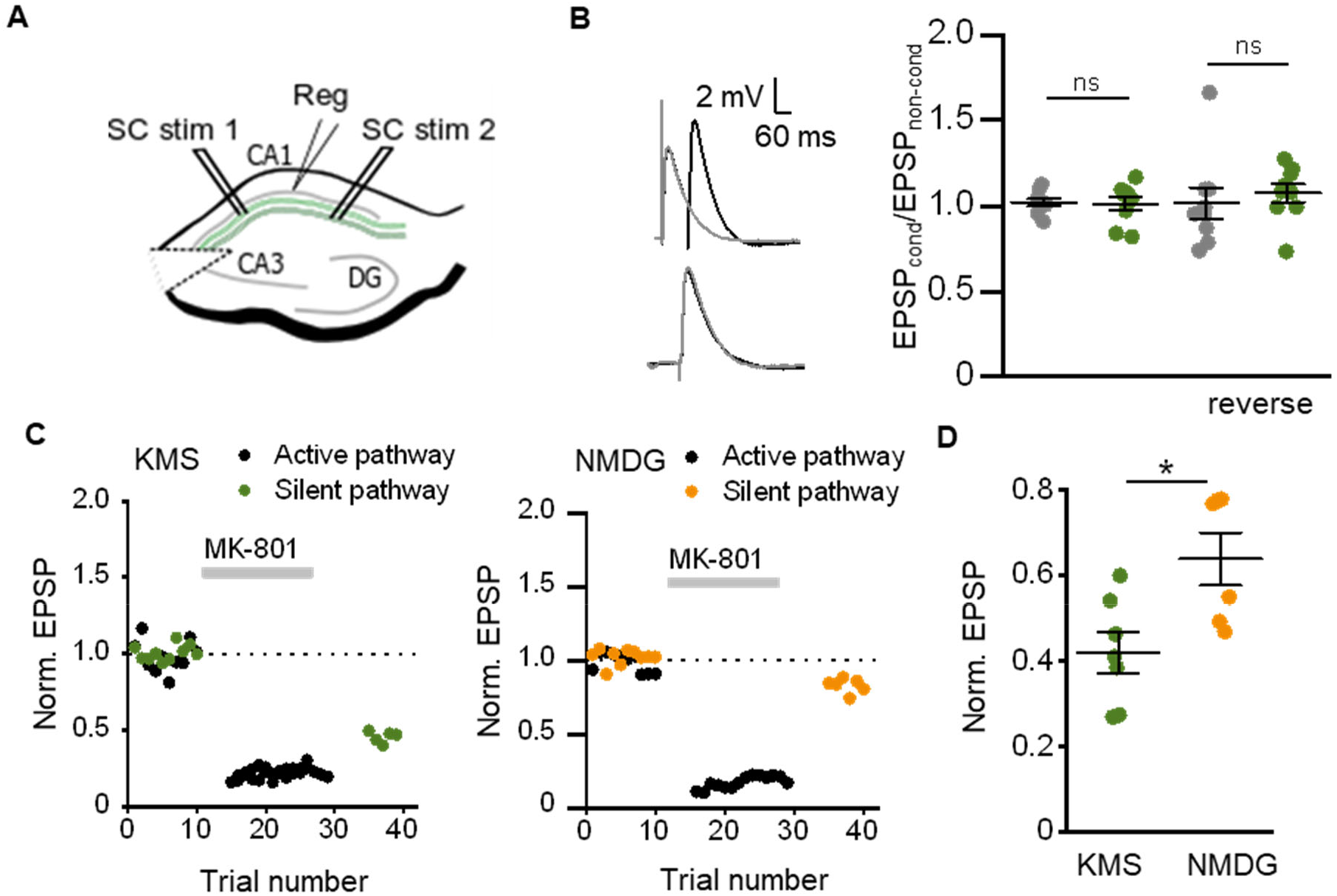
Postsynaptic K^+^ efflux enhances intersynaptic crosstalk. **A.** A schematic showing the position of two stimulating (SC stim 1 and SC stim 2) and one recording (Reg) electrodes. Stimulating electrodes were placed to activate two parallel pathways of Schaffer collaterals. EPSPs were recorded from CA1 pyramidal neurons. **B.** The test on pathway independence. *Left*, sample traces of conditioned EPSP (EPSP in one pathway was preceded by EPSP in the other pathway) and unconditioned EPSP. *Right*, a ratio of conditional EPSP versus non‐conditional EPSP **C.** Examples of two pathway recordings with KMS‐ (*left*) and NMDG‐based (*right*) internal solutions. Grey bar ‐ application of MK‐801, activity‐dependent channel blocker of NMDARs. Replacement of intracellular K^+^ for NMDG reduced the blockade of the silent pathway during stimulation of the active pathway. This points to reduced glutamate spillover. **D.** The summary plot showing EPSP reduction in the silent pathway. The EPSPs were normalized to their baseline amplitude. The reduction was significantly larger when the postsynaptic cell was filled with K^+^. The data are presented as mean ± SEM. ns p > 0.05, *p < 0.05, two‐sample *t*‐test.

## Discussion

In the course of excitatory synaptic transmission, astrocytes clear released glutamate and excess of K^+^ from the extracellular space (Verkhratsky and Nedergaard, 2018). We observed an activity‐dependent increase in the amplitude and τ_decay_ of synaptically‐induced I_GluT_ in hippocampal astrocytes. The increase of the amplitude of I_GluT_ is consistent with the facilitation of presynaptic release probability mediated by K^+^ accumulation in the synaptic cleft, as reported earlier (Contini et al., 2017; Shih et al., 2013). Indeed, this increase was suppressed by the blockade or genetic deletion of the postsynaptic of NMDARs, which serve as a major source of K^+^ efflux during synaptic transmission. Similarly, the increase of τ_decay_ was abolished in these tests. The result is consistent with the early report suggesting that glutamate transporters are not overwhelmed during HFS upon blockade of ionotropic postsynaptic receptors (Diamond and Jahr, 2000). Thus, glutamate transporters can effectively clear glutamate in hippocampal CA1 unless local glutamate translocation is partly suppressed by K^+^ efflux through postsynaptic NMDARs. Depolarization of perisynaptic astrocytic leaflets or a decrease of the transmembrane K^+^ gradient, or both, can potentially underpin reduced glutamate uptake (Grewer et al., 2008). However, we found that K^+^ elevation alone does not affect I_GluT_, while astrocyte depolarization increases its τ_decay_. How much K^+^ accumulation during synaptic transmission can depolarize leaflets cannot be directly measured because these processes are beyond optical resolution for light‐microscopy voltage imaging, nor can it be accessed with electrode‐based techniques. Previous simulation studies suggest that K^+^ can rise up to 5 mM in the synaptic cleft during a single EPSC (Shih et al., 2013). However, the K^+^ concentration drops rapidly outside of synaptic cleft, and its effect on the astrocyte may be strongly attenuated. Nevertheless, space and time‐averaged extracellular [K^+^]_o_ during HFS can increase by several millimoles (Durand Dominique et al., 2010). In such cases, the local perisynaptic K^+^ elevation should be several folds higher. Although it was not technically feasible to probe the effects of K^+^ hotspots on the microscopic scale, we found that elevation of K^+^ to 7.5 mM depolarizes the astrocyte from −83.5 ± 1.19 mV to −65.2 ± 3.41 mV, which significantly increases τ_decay_ of uI_GluT_.

Whilst changes in I_GluT_ reflect glutamate translocation across the cell membrane, the extracellular glutamate concentration could follow a different time course. Reduced transporter turnover may increase the number of extracellular glutamate transporter binding sites available for glutamate and thus boost the transporter buffering capacity. Increased buffering could slow down glutamate diffusion and prolong the glutamate transient tail. Indeed, imaging with glutamate sensor iGluSnFR revealed a decrease in τ_decay_ of glutamate transient when K^+^ in the postsynaptic cell was replaced with NMDG. Consistent with a longer glutamate dwell‐time, the activity‐dependent prolongation of EPSPs was observed only when the postsynaptic cell contained a physiological range concentration of K^+^, but not when K^+^ was replaced with NMDG. However, an increase in local glutamate dwell‐time does not reveal the efficiency of glutamate spillover. The latter may be enhanced with reduced glutamate uptake and reduced with increased glutamate buffering, or indeed under withdrawal of perisynaptic leaflets (Henneberger et al., 2020). We employed a two‐pathway experiment to demonstrate that glutamate spillover also depends on postsynaptic K^+^ efflux.

Our findings suggest that blockade of postsynaptic NMDARs reduces activity‐dependent facilitation of glutamate release and spillover. However, an alternative explanation is that blocking postsynaptic receptors in the entire slice suppresses recurrent excitation of CA3 pyramidal neurons and hence activity‐dependent recruitment of additional CA3‐>CA1 fibers. While this could explain the NMDAR‐dependent facilitation of I_GluT_, it could not explain the documented prolongation of I_GluT_ decay time. Moreover, the CA1‐GluN1‐KO mice showed reduced activity‐dependent facilitation and the reduced prolongation of I_GluT_. Replacing intracellular K^+^ for NMDG similarly reduced the activity‐dependent facilitation and prolongation of EPSPs in pyramidal neurons. Since CA3 pyramidal neurons were not affected in either experiment, these results effectively rule out polysynaptic effects.

The source of K^+^ released by the postsynaptic neuron is another crucial issue. In addition to their own Ca^2+^ permeability, NMDARs are linked to Ca^2+^ dependent K^+^ channels that can potentially contribute to K^+^ efflux (Ngo‐Anh et al., 2005; Shah and Haylett, 2002; Zhang et al., 2018). However, extracellular Ca^2+^ removal does not reduce I_K_ in astrocytes induced by activation of neuronal NMDARs with glutamate puff (Shih et al., 2013). Hence, a significant contribution of Ca^2+^ ‐dependent K^+^ channels to K^+^ efflux during synaptic transmission is unlikely. Another possibility is the activation of voltage‐gated K^+^ channels during EPSPs. However, blockade of NMDARs reduces field EPSPs to a much lesser extent than it reduces I_K_ (Shih et al., 2013). Thus, no major contribution of voltage‐gated K^+^ channels to K^+^ efflux during synaptic transmission has been observed.

Another critical question is to what extent K^+^ released at an individual synapse affects glutamate uptake in perisynaptic astrocytic leaflets. Biophysical models predict that K^+^ concentration rapidly drops with distance from the cleft, with no effect on astrocytic processes at nearest‐neighbor synapses (Shih et al., 2013). The low input resistance combined with a large membrane area of astroglia severely limits the spread of current‐triggered membrane depolarization in astrocytes. Therefore, increasing the number of activated synapses should not affect the kinetics of glutamate uptake unless their K^+^ hotspots become overlapped. The latter scenario could occur during synchronous activation of multiple neighboring synapses or during epileptic bursts. Such events do produce wide‐spread elevations of extracellular K^+^ affecting large astrocyte territories.

In summary, we conclude that glutamate spillover is prevented by efficient glutamate uptake unless astroglial transporters are downregulated or withdrawn from the immediate synaptic environment. Indeed, early reports demonstrated that lowering recording temperature to decrease glutamate uptake efficiently promotes glutamate spillover (Asztely et al., 1997; Kullmann and Asztely, 1998). Also, early findings highlighted the role of NMDARs to detect spillover while attributing this fact to higher glutamate affinity of these receptors (Diamond, 2001; Rusakov and Kullmann, 1998).

Our finding demonstrates that K^+^ efflux through postsynaptic NMDARs depolarizes astrocytic leaflets and thus reduces local glutamate uptake. What could be the physiological role of this phenomenon? In hippocampal circuitry, enhanced glutamate spillover has been associated with co‐operative action of dendritic NMDARs, such as receptor ‘priming’ (Arnth‐Jensen et al., 2002; Hires et al., 2008). It has been found to underlie functional inter‐synaptic crosstalk (Arnth‐Jensen et al., 2002; Lozovaya et al., 1999; Scimemi et al., 2004), also contributing significantly to heterosynaptic plasticity (Vogt and Nicoll, 1999), and activation of metabotropic glutamate receptors outside the synaptic cleft (Min et al., 1998; Semyanov and Kullmann, 2000).

Recent reports suggest that a decrease in glutamate uptake can shift the sign of synaptic plasticity, reducing long‐term potentiation (LTP) and promote long‐term depression (LTD) (Valtcheva and Venance, 2019). Rate‐based LTP induction in one set of synapses requires NMDARs activation and thus should lead to K^+^ hotspots in the perisynaptic space. This K^+^ should, in turn, depolarize astrocytic leaflets and downregulate glutamate uptake. However, leaflets are ‘shared’ between neighboring synapses (Gavrilov et al., 2018). We thus speculate that LTP in one set of synapses should suppress LTP and facilitate LTD in their neighbors if they are activated immediately after while astrocyte is depolarized.

LTP itself changes the synapse's ability to release K^+^ to the extrasynaptic space (Ge and Duan, 2007). After the LTP, the number of AMPARs is increased. AMPARs do not contribute significantly to K^+^ efflux due to their fast inactivation but can facilitate activation of NMDARs by removing their voltage‐dependent block (Shih et al., 2013). Therefore, LTP not only increases the quantal efficiency of the synapse but promotes K^+^‐dependent facilitation of glutamate release and spillover at this synapse, potentially exceeding the effect of temporal perisynaptic astrocytic leaflet withdrawal after LTP induction (Henneberger et al., 2020). K^+^‐ dependent facilitation of presynaptic glutamate release after LTP could be linked to the putative perisynaptic mechanism of LTP (Kullmann, 2012).

Our finding emphasizes the physiological importance of changes in ionic concentrations in the synaptic cleft and perisynaptic space. Since the volumes of these spaces are minimal, the concentration changes could be significant. Accumulation of K^+^ in the synaptic cleft is paralleled by local Ca^2+^ depletion (Rusakov and Fine, 2003), which is also sensed by astrocytes (Torres et al., 2012) and might affect release efficacy in the opposite direction to that of excess K^+^. How the astrocyte mediated K^+^ buffering affects the time‐course of perisynaptic K^+^ elevation and how far K^+^ can diffuse in the extracellular space remains to be established.

## Materials and Methods

### Hippocampal Slice Preparation

Animal procedures were carried out under the oversight of the UK Home Office (as per the European Commission Directive 86/609/EEC and the UK Animals (Scientific Procedures) Act, 1986) and by institutional guidelines. Young C57BL/6 mice (3–4 weeks of age) male were anesthetized using isoflurane and decapitated. The brain was exposed, chilled with an ice‐cold solution containing (in mM): sucrose 75, NaCl 87, KCl 2.5, CaCl_2_ 0.5, NaH_2_PO_4_ 1.25, MgCl_2_ 7, NaHCO_3_ 25, and D‐glucose 25. Hippocampi from both hemispheres were isolated and placed in an agar block. Transverse slices (350 μm) were cut with a vibrating microtome (LeicaVT1200S) and left to recover for 20 min in the same solution at 34°C. Then slices were incubated at 34°C in a solution containing (in mM): NaCl 119, KCl 2.5, NaH_2_PO_4_ 1.25, MgSO_4_ 1.3, CaCl_2_ 2.5, NaHCO_3_ 25, and D‐glucose 11. For experiments with intracellular blockade of K_ir_, the solution was supplemented with 100 μM BaCl_2_. After that, slices were transferred to the recording chamber and continuously perfused at 34°C with the same solution. All solutions were saturated with 95% O2 and 5% CO2. Osmolarity was adjusted to 298 ± 3 mOsM.

### AAV Transduction

For viral gene delivery of AAV2/1h.Synap.SF‐iGluSnFR‐A184V (Penn Vector Core, PA, USA), pups, male and female (P0‐P1), were prepared for aseptic surgery. To ensure proper delivery, intracerebroventricular (ICV) injections were carried out after an adequate visualization of the targeted area (Kim et al., 2014). Viral particles were injected in a volume 2.5 μl/hemisphere (totaling 5 × 10^9^ genomic copies), using a glass Hamilton microsyringe at a rate not exceeding of 0.2 μl/s, 2 mm deep, perpendicular to the skull surface, guided to a location approximately 0.25 mm lateral to the sagittal suture and 0.50–0.75 mm rostral to the neonatal coronary suture. Once delivery was completed, the microsyringe was left in place for 20–30 s before being retracted. Pups (while away from mothers) were continuously maintained in a warm environment to eliminate the risk of hypothermia in neonates. After animals received AAV injections, they were returned to the mother in their home cage. Pups were systematically kept as a group of litters. Every animal was closely monitored for signs of hypothermia following the procedure and for days thereafter to ensure that no detrimental side effects appear. For the transduction of glutamate sensors in vivo, there were three‐ to four‐ weeks to suffice.

### Electrophysiology

Whole‐cell recordings from *stratum radiatum* astrocytes were obtained using patch electrodes filled with a potassium methane sulfonate‐based solution (KMS) containing (in mM): CH_3_KO_3_S 135, HEPES 10, MgCl_2_ 4, disodium phosphocreatine 10, Na_2_ATP 4, NaGTP 0.4 (pH adjusted to 7.2 with KOH; osmolarity to 290 ± 3 mOsM) with a resistance of 3–5 MΩ. 50 μM Alexa Fluor 594 was added to the intracellular solution as a morphological marker. Astrocytes were identified by small soma size (about 10 μm), low resting membrane potential (−84.0 ± 0.5 mV, *n =* 16), and low input resistance (16.3 ± 1.4 MΩ, *n =* 16). Passive cell properties were confirmed by linear I–V characteristics (Fig. 1B). Synaptic responses were evoked by single and burst stimulation (5 stimuli at 50 Hz) of Schaffer Collaterals (SC) with a bipolar electrode (FHC, Bowdoin, USA). The stimulating electrode was placed in the *stratum radiatum* more than 200 μm from the recording site. Signals were amplified with the Multiclamp 700B amplifier (Molecular Devices), filtered at 2 kHz, recorded and digitized at 4– 10 kHz with the NI PCI‐6221 card (National Instruments).

Whole‐cell recordings of CA1 pyramidal neurons were obtained using patch electrodes filled with a solution containing (in mM): CH_3_KO_3_S 130, NaCl 8, HEPES 10, disodium phosphocreatine 10, Na_2_GTP 0.4, MgATP 4, and 3 mM Na‐ascorbate (pH adjusted to 7.2 with KOH; osmolarity to 290 ± 3 mOsM). To eliminate postsynaptic K^+^ efflux, CH_3_KO_3_S was replaced with N‐methyl‐D‐glucamine CH_3_SO_3_ (NMDG).

### Glutamate Uncaging

4‐methoxy‐7‐nitroindolinyl‐caged L‐glutamate (10 mM, MNI‐glutamate) was applied to the perfusion solution. Glutamate uncaging was carried out using mode‐locked tunable 690– 1020 nm laser Mai‐Tai (Spectra‐Physics, USA) in a “point scan” mode for 5 ms at 720 nm with the FV1000‐MPE system.

### iGluSnFR Imaging

Femtonics Femto2D‐FLIM imaging system was used for two‐photon imaging, integrated with two femtosecond pulse lasers MaiTai (SpectraPhysics‐Newport) with independent shutter and intensity control and patch‐clamp electrophysiology system (Femtonics, Budapest). Patch pipettes were pulled from borosilicate–standard wall filament glass (G150F‐ 4; Warner Instruments, CT, USA) with 4–5 MΩ resistance. CA1 pyramidal neurons expressing iGluSnFR sensor were patch‐clamped with either KMS‐ or NMDG‐based internal solution, supplemented with the morphological tracer dye Alexa Fluor 594 (50 μM). The Alexa Fluor 594 channel was used to identify a region of interest and recorded along with the iGluSnFR signal. After at least 45 min required for the dye diffusion and equilibration in the dendritic arbor, glutamate imaging from individual spines was carried out using an adaptation of the previously described method (Jensen et al., 2017).For the fast imaging of AP‐mediated glutamate transients point scans were performed over the dendritic spines and scanned at a sampling frequency of 500 Hz.

### Astrocyte simulations

Simulations were carried out using a detailed 3D biophysical model of a (statistically) reconstructed CA1 astrocyte using a NEURON‐compatible model builder ASTRO (www.neuroalgebra.com and https://github.com/LeonidSavtchenko/Astro), as outlined in detail earlier (Savtchenko et al., 2018). In brief, ‘average’ astrocyte morphology was obtained from a sample of protoplasmic astrocyte in hippocampal *stratum radiatum* (~4 week‐old rats) using the systematic procedures as follows. First, generating soma and primary (optically discernible) branches to match sample‐average numbers and dimensions; second, adding nanoscopic branches that have statistically generated dimensions matching experimental EM data (size distributions); third, adjusting biophysical properties of nano‐branches to match biophysics of reconstructed 3D EM processes; fourth, adjusting the numbers and density of nano‐branches to match the statistics of tissue volume fraction and surface‐to‐volume ratios obtained empirically from 3D EM data; fifth, distributing biophysical membrane mechanisms across the model cell membrane, including K_ir_4.1 channels and GLT‐1 transporters in accord with the experimental recordings. A full description of the model and its implementation, for either desktop‐ or cloud‐computing, are available from www.neuralgebra.com and links therein.

The model was populated with K_ir_4.1 channel, with the kinetics in accord with (Sibille et al., 2015), and membrane unit conductance of *GKir* =0.1 mS cm^−2^.

The kinetic is described by the equation:

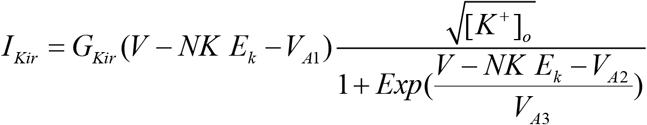

Where *V*_*A1*_ = ‐14.83 mV, *V*_*A2*_ = ‐105.82 mV, *V*_*A3*_ = 19.23 mV, *NK* = 0.81, *E*_*k*_ is the Nernst astrocyte K^+^ potential, *V* is the astrocyte membrane potential, [*K^+^]*_*0*_ is the extracellular *K*^*+*^ concentration and *V*_*A1*_ (an equilibrium parameter, which sets I_Kir_ current to 0 at −80 mV), *NK,V*_*A2*_ and *V*_*A3*_ are constant parameters calibrated by the I‐V curve.

The leak passive current *I*_*pas*_ = *G*_*pas*_(*V* − *E*_*pas*_) was added to stabilize the astrocyte membrane potential at *E*_*pas*_ =−85 mV, *G*_*pas*_=0.001 mS/cm^2^.

GLT‐1 kinetics was modelled in accord with (Bergles et al., 2002; Zhang et al., 2007), with the surface density of 10^4^ μm^‐2^ as estimated earlier (Lehre and Danbolt, 1998).

The kinetics of glutamate transporters was determined by a simplified scheme of 6 states:

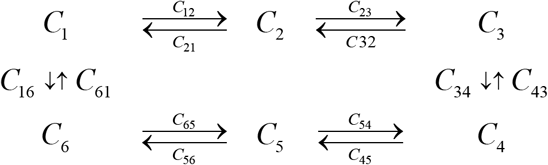

Where C_12_=[Glu]_o_ k_12_ u(v,−0.1), C_21_=k_21_, C_23_=[Na^+^]_o_ k_23_ u(v,0.5), C_32_=k_32_, C_34_=k_34_ u(V,0.4), C_43_=k_43_, C_45_=k_45_, C_54_=k_54_ [Glu]_in_, C_56_=k_56_ u(v,0.6), C_65_=k_65_ [Na^+^]_in_, C_61_=[K^+^]_in_ k_61_, C_16_=k_16_ u(V,0.6) [K^+^]_o_.

And function u(V,P)= exp(P V/53.4), where V is astrocyte membrane voltage, and P is a charge translation between states C_i_ ‐> C_j_.

The value of kinetic constant were : k_12_=20 /mM /ms, k_21_=0.1 /ms, k_23_=0.015 /mM /ms, k_32_=0.5 /ms, k34=0.2 /ms, k_43_=0.6 /ms, k_45_=4 /ms, k_54_=10 /mM/ms, k_56_=1 /ms, k_65_=0.1/mM/ms, k_16_=0.0016 /mM/ms, k_61_=2 10^‐4^ /mM /ms.

The glutamate transporter current (Zhang et al., 2007) was calculated according to the following equation :

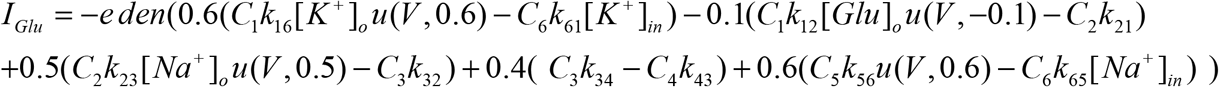

Where e is charge on an electron charge 1.6×10^−19^ (coulombs) and den is a density of transporters

Initial ion concentrations:

[Na^+^]_in_=15 mM, [Na^+^]_o_=150 mM, [K^+^]_in_=120 mM, [K^+^]_o_=3 mM, [Glu]_in_=0.3 mM, Diffusion of intracellular potassium is described by the equation built into the Neuron : ~ ka << (−I_Kir4.1_/(Fπd)), where F= 96485 C/M is Faraday constant, d is a local diameter with potassium diffusion coefficient D_K_=0.6 mm/ms^2^.

To fit the experimental observation, we fit the kinetic scheme and modified the sensitivity of transporter to the change of voltage when binding [K]o, C16=k16 u(V,0.6) [K+]o from 0.6 to 0.1.

Extracellular [K^+^]_o_ elevation from 2.5 mM up to 5 mM was simulated across a 20 μm spherical area within the astrocyte arbor: while diffusing away, it also activated Kir4.1 homogeneously within the area, prompting K^+^ entry into the astrocyte (peak current density ~0.01 mA cm^‐2^). The ensuing local increase in intracellular K^+^ concentration (from 110 to ~110.5 mM) dissipated over several seconds after extracellular K^+^ concentration returned to the baseline value of 2.5 mM. Extracellular glutamate rise (peak 0.1 mM, 2 ms) was simulated within a 3 μm sphere inside the astrocyte arbor.

The extracellular concentration of glutamate was calculated as a conditions IGlu=0 at [K]o=5 [Glu]o=8.5*10^−5^ mM and [K]=2.5 mM [Glu]o=3.57*10^−6^ mM.

### Drugs and Chemicals

All drugs were made from stock solutions kept frozen at −20°C in 100–200 ml 1,000×aliquots. DL‐2‐amino‐5‐phosphonovaleric acid (D‐APV), 2,3‐dioxo‐6‐nitro‐7‐sulfamoyl‐benzo[f]quinoxaline (NBQX), DL‐threo‐β‐benzyloxyaspartic acid (DL‐TBOA), [5R,10S]‐[+]‐5‐ methyl‐10,11‐dihydro‐5H‐dibenzo[a,d]cyclohepten‐5,10‐imine (MK‐801),picrotoxin, and MNI‐glutamate were purchased from Tocris Cookson (Bristol, UK). Chemicals for solutions were from Sigma‐Aldrich (St. Louis, USA). Alexa Fluor 594 was obtained from Invitrogen (Carlsbad, USA).

### Statistical Analysis

Electrophysiological data were analyzed with WinWCP and Clampfit (Axon Instruments Inc.; Union City, USA). Imaging data were analyzed using MES software (Femtonics, Budapest), ImageJ (a public domain Java image processing program by Wayne Rasband), and traces expressed as ΔF/F. Statistical analysis was performed using Excel (Microsoft, US), Origin 8 (Origin Lab Corp, Northampton, USA). All data are presented as the mean ± SEM, and the statistical difference between means was estimated with the unpaired *t*‐test and repeated measurements two‐way ANOVA as stated in the text.

## Supplementary figure

**Figure S1.**
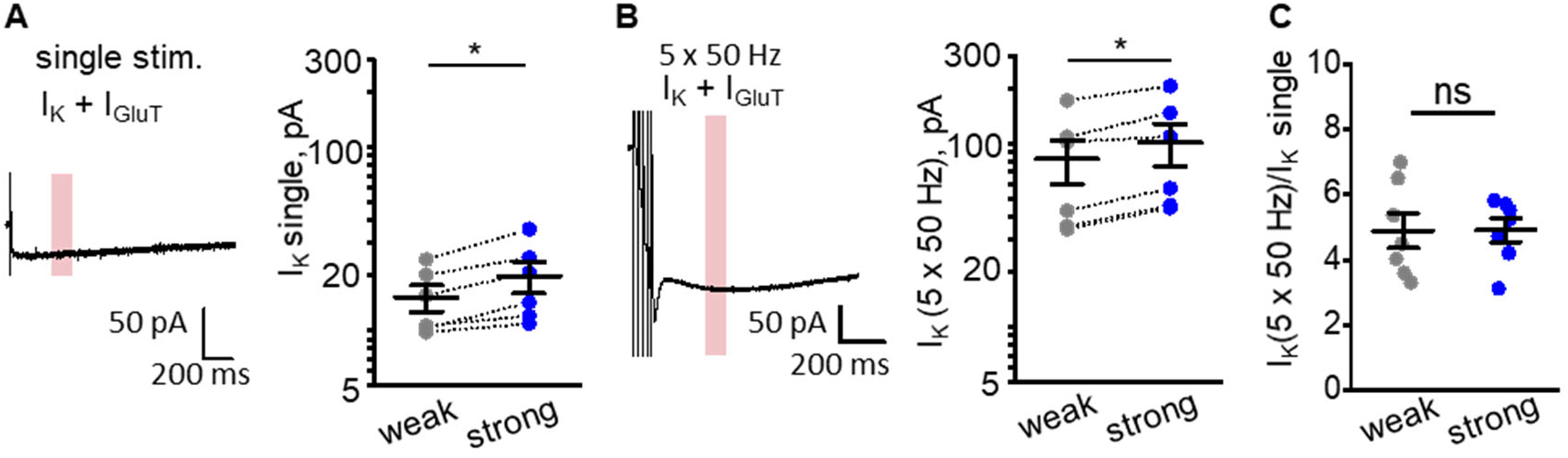
Astrocytic K^+^ current scales with increases in stimulus strength but shows the unchanged use‐dependent increase during bursts. **A.** Trace of characteristic *I*_*K*_+*I*_*GluT*_ current in response to a single stimulus applied to Shaffer collaterals; pink bar (200 ms after the last peak): current measurement window. Graph: statistical summary showing *I*_*K*_ values during weak and strong stimuli. **B.** Experiments as in (A), but with a 5 × 50 Hz stimulus burst; notation as in (A). **C.** The summary graphs showing an *I*_*K*_ increase during 5 × 50 Hz bursts relative to the single stimulus *I*_*k*_. The data are presented as mean ± SEM. ns p > 0.05 and *p < 0.05, two‐sample t‐test.

